# Novel Mechanisms of Strigolactone-Induced DWARF14 Degradation in *Arabidopsis thaliana*

**DOI:** 10.1101/2024.01.10.574527

**Authors:** Elena Sánchez Martín-Fontecha, Francesca Cardinale, Marco Bürger, Cristina Prandi, Pilar Cubas

## Abstract

Strigolactones (SLs) are carotenoid-derived compounds that regulate various aspects of plant development, physiological responses and plant interactions with microorganisms. In angiosperms, the SL receptor is the α/β hydrolase D14 that, upon SL binding, undergoes conformational changes, triggers SL-dependent responses and hydrolyses SLs. Arabidopsis SL signalling involves the formation of a complex between SL-bound D14, the E3-ubiquitin ligase SCF^MAX2^ and the transcriptional corepressors SMXL6/7/8 (SMXLs), which become ubiquitinated and degraded by the proteasome. However, the sequence of events that result in SL signalling and the requirement of SL hydrolysis for this process remain unclear. In addition, SL destabilises the D14 receptor. The biological significance of SL-induced D14 degradation is unclear, although it has been proposed to create a negative feedback loop in SL signalling. The current model proposes that D14 degradation occurs after SMXLs ubiquitination and proteolysis, via the same E3-ubiquitin ligase that targets the repressors.

In this work we quantitatively studied the degradation dynamics of Arabidopsis D14 in response to SLs *in planta*. For this, we conducted fluorescence and luminescence assays to monitor D14 stability dynamics upon SL treatments, in transgenic lines expressing *D14* fused to *GREEN FLUORESCENT PROTEIN* (*GFP*) or *LUCIFERASE* (*LUC*), in wild-type and SL-signalling mutant backgrounds. Mutant D14 proteins predicted to be non-functional for SL signalling were also examined, and their capability to bind SLs *in vitro* was studied using Differential Scanning Fluorimetry (DSF). Finally, we used a non-hydrolysable SL to test the requirements of SL hydrolysis for D14 and SMXL7 degradation. Our research revealed that SL-induced D14 degradation may occur in the absence of SCF^MAX2^ and/or SMXLs by a proteasome-independent mechanism. Additionally, we observed conditions in which the efficiency of SL-induced degradation of D14 is not aligned with that of SMXL7 degradation. Finally, our results indicate that the hydrolysis of SLs is not a prerequisite to trigger either D14 or SMXL7 degradation. These findings suggest the existence of a regulatory mechanism governing D14 degradation more complex than anticipated, and provide novel insights into the dynamics of SL signalling in Arabidopsis.

## Introduction

Strigolactones (SLs) are a class of carotenoid-derived compounds (Al-Babili and Bouwmeester, 2015) known to act as exuded and endogenous signals in both beneficial and detrimental interactions with (micro)organisms in the rhizosphere (Akiyama *et al*., 2005; Besserer *et al*., 2006). They also play crucial roles in the control of plant development and growth: they are inhibitors of bud outgrowth and shoot branching, and regulators of internode elongation, height, stem secondary growth, leaf development and senescence, reproduction and of root architecture (reviewed in Rameau *et al*., 2019; Dun *et al*., 2023; Guercio *et al*., 2023). SLs are mediators of physiological and morphological responses to water and nutrient deprivation, including the ability to form mycorrhizae and nodules, and are important for full antioxidant capacity in response to stress (reviewed by Trasoletti *et al*., 2022; Lanfranco *et al*., 2018). Their perception is repressed by sugars and citrate, that may act as proxies for the plant nutritional status (Tal *et al*., 2022; reviewed by Barbier *et al*., 2023). Thus, they are thought to integrate environmental signals with plant plasticity and physiological acclimation.

Structurally, SLs are tricyclic lactones (ABC tricyclic system) connected to a butenolide (D-ring) through an enol-ether bond (reviewed by Dun *et al*., 2023). There are two key stereochemical centres in these molecules, one at the junction of the B and C rings, and the other at the junction of the D-ring. The first stereocentre is not critical in defining SL activity, but does divide SLs into two subclasses, the 5-deoxystrigol (5DS) and 4-deoxyorobanchol (4DO) types. Naturally occurring plant SLs display a 2’R configuration of the D-ring (reviewed in Yoneyama *et al*., 2018), which is required for their biological activity.

The signalling mechanism of SLs is based on a hormone-induced proteolysis pathway similar to that of other plant hormones such as auxins, gibberellins and jasmonate. It involves a hormone receptor, and a Skp1-Cullin-F-box (SCF) E3-ubiquitin ligase complex that targets specific protein substrates for polyubiquitination and subsequent degradation by the 26S proteasome (Blázquez *et al*., 2020). In angiosperms, the SL receptor is the α/β hydrolase D14, unusual insofar as it can hydrolyse its ligand, albeit at a low turnover rate. D14 contains a hydrophobic binding pocket that can directly bind to the SL molecule (reviewed in Barbier *et al.,* 2023; Guercio *et al.,* 2023). The protein active site is located at the bottom of the SL binding pocket and is formed by a conserved catalytic triad, serine-histidine-aspartic acid (S97-H247-D218 in Arabidopsis).

SL signalling is initiated with the binding of SLs to D14. The nucleophilic attack of C5’of SL by the catalytic serine is followed by several states involving intermediates that may affect the enzymatic activity of D14 (reviewed in Guercio *et al.,* 2023). SL signalling also requires the formation of a complex between SL-D14, the E3-ubiquitin ligase complex SCF^MAX2/D3^, which confers the substrate specificity by the F-box protein MORE AXILLARY GROWTH2 (MAX2) or D3 (in Arabidopsis and rice respectively, Ryun Woo *et al*., 2001; Stirnberg *et al*., 2002, 2007; Ishikawa *et al*., 2005; Johnson *et al*., 2006), and the transcriptional corepressors SUPPRESSOR OF MAX2-1-LIKE (SMXL)6, SMXL7 and SMXL8 (hereafter SMXLs) or DWARF53 (D53) in Arabidopsis or rice respectively (Jiang *et al*., 2013; Zhou *et al*., 2013; Soundappan *et al*., 2015; Wang *et al*., 2015; Liang *et al*., 2016). SMXLs/D53 are targets for polyubiquitination by SCF^MAX2/D3^ and proteasomal degradation, which elicits SL-dependent gene expression (Jiang *et al*., 2013; Zhou *et al*., 2013; Soundappan *et al*., 2015; Wang *et al*., 2015; Liang *et al*., 2016). To date, the sequence of events leading to complex formation and whether SL hydrolysis is required for this process are yet unclear.

The formation of the complex and the subsequent SL signalling requires SL-induced conformational changes in D14. One model, based on structural data, proposes that SL hydrolysis is essential for these conformational changes (Yao *et al.,* 2016; Saint Germain *et al*., 2016). A second model, based on time-course analyses of SL binding and hydrolysis and genetic data, proposes that the intact SL molecule triggers complex formation and signalling. This model is further supported by the observation that the recruitment of D3 and D53 requires a pre-hydrolysis state of D14-SL (Shabek *et al.,* 2018; Seto *et al.,* 2019). In this model, hydrolysis of SLs would serve to deactivate the bioactive form of the hormone (Seto *et al.,* 2019). Nevertheless, the exact series of events, or whether ligand hydrolysis is indeed required for SL signalling, is not settled yet. Furthermore, D14 may not be a single turnover enzyme, as the interaction between the hydrolysis intermediate and D14 seems to be reversible. This implies that the receptor could be reused and participate in several SL perception/deactivation cycles (Shabek *et al*., 2018).

After SL-induced ubiquitination and degradation of the SMXLs/D53, the SL receptor D14 is itself destabilised in the presence of SLs. This suggests that SLs may promote a negative feedback loop within the SL signalling cascade, by limiting the availability of receptors and thus modulating the sensitivity to SLs (Chevalier *et al*., 2014). Evidence for the phenomenon of SL-induced destabilisation of D14 has been found in Arabidopsis and rice. Arabidopsis transgenic lines expressing translational fusions of *D14* and *GREEN FLUORESCENT PROTEIN* (*GFP*) or *β-GLUCURONIDASE* (*GUS*) coding sequences (CDS) display a significant decrease in D14:GFP and D14:GUS protein levels, respectively, following treatments with SLs (Chevalier *et al*., 2014). Cell-free degradation assays using Arabidopsis plant extracts and a purified D14:HA protein show reduced D14:HA protein abundance when exposed to SLs (Tal *et al*., 2022). Rice *D14:GFP*- and *HA:D14*-expressing calli display D14:GFP and HA:D14 destabilisation after addition of SLs (Hu *et al.,* 2017; Patil *et al.,* 2022).

The initially proposed model for D14 degradation hypothesises that, following ubiquitination and proteasomal removal of SMXLs/D53, D14 becomes accessible to ubiquitination and proteasomal degradation by the same SCF^MAX2/D3^ complex (Chevalier *et al*., 2014; Hu *et al*., 2017; Tal *et al*., 2022). Several observations support this scenario: SL treatments induce ubiquitination of D14:GFP in rice (Hu *et al*., 2017); degradation of Arabidopsis and rice D14:GFP can be reduced by proteasome inhibitors (Chevalier *et al*., 2014; Hu *et al*., 2017); destabilisation of D14:GFP is decreased in Arabidopsis *max2* and rice *d3* mutants (Chevalier *et al*., 2014; Hu *et al*., 2017). Moreover, rice SL-insensitive *d53* mutants render D14:GFP resistant to SL-induced degradation, which led to propose that D14 and D53 degradation are coupled, and that both are strongly associated to the SL signalling status (Hu *et al*., 2017). Finally, D14 mutant analyses suggest that SL hydrolysis is essential for D14 degradation (Hu *et al*., 2017).

This general model is derived from the amalgamation of a wide array of heterogeneous sources, most of them involving callus cultures and cell-free assays performed mainly in rice. Therefore, it remains largely untested *in planta* whether MAX2 and SMXLs are indispensable for the full extent of SL-induced D14 degradation in Arabidopsis. Moreover, the biological significance of the SL-induced D14 degradation and its relationship with SL signalling remains unclear.

In this work we systematically studied, quantitatively and *in planta,* the degradation dynamics of the Arabidopsis D14 protein (AtD14, hereafter D14) in response to SLs. With this aim we generated transgenic lines constitutively expressing the *D14* CDS fused to that of the *GFP* or *LUCIFERASE (LUC)*, in wild-type and mutant backgrounds affected in the SL signalling machinery. We then measured D14:GFP fluorescence signal (using time-lapse fluorescence quantitative microphotography) or D14:LUC activity (using luminescence assays) in plants treated with the synthetic SL analogue GR24^5DS^. We performed similar assays with D14 proteins with mutations in residues thought to be critical for D14 activity or for interaction with other components of the SL signalling machinery. The ability of these mutants to bind SLs was tested by Differential Scanning Fluorimetry (DSF). Additionally, we used synthetic, non-hydrolysable SL derivatives to assess the requirement for SL hydrolysis during this process.

Our results reveal novel and unexpected features of D14 degradation in Arabidopsis that are inconsistent with the current models being exclusive. We show that SL hydrolysis upon binding to D14 is not essential for the degradation of the receptor. We also found indications of a proteasome-independent mechanism of D14 degradation and provided compelling evidence that neither MAX2 nor the SMXLs are strictly essential for D14 degradation, although they do contribute to the process. Finally, we report cases in which the efficiency of SL-induced D14 degradation is largely divergent from that of SMXL7. All these results indicate that, in addition to the SCF^MAX2^/SMXLs-associated mode of D14 degradation, there are alternative pathways to modulate D14 levels in the presence of SLs.

## Material and Methods

### Plant material

Wild-type *Arabidopsis thaliana* plants of the Columbia-0 (Col-0) ecotype were used. The *d14-1* mutant (Arite *et al*., 2009; Waters *et al*., 2012), was obtained from the Nottingham Arabidopsis Stock Centre (NASC ID:N913109). The *max2-1* mutant (Stirnberg *et al*., 2002) was provided by Dr. Ottoline Leyser. T-DNA insertional lines *smxl6-4 smxl7-3 smxl8-1* (*s678*) and *smxl6-4 smxl7-3 smxl8-1 max2-1* (*s678m*) (Liang *et al*., 2016) were provided by Dr. Tom Bennett. Experiments were performed in homozygous lines, except for the SMXL7:LUC degradation assays in *35S:D14^P169L^:GFP;d14-1 and 35S:D14^G158E^:GFP;d14-1* background that were performed in F1 hemizygous lines from crosses of these lines with *UB:SMXL7:LUC;d14-1*.

### Growth conditions

Arabidopsis seeds were sown in trays containing a mix of commercial soil and vermiculite (3:1 proportion), stratified in darkness 2-3 days at 4 °C and grown in long day conditions (16 h light/8 h dark). Humidity rate was 60%, temperature 22 °C and white light (photosynthetically active radiation, 100 μmol·m^−2^s^−1^) was provided by cool-white 20-W F20T12/CW tubes (Phillips). For *in vitro* studies, seeds were surface-sterilised by 7 min incubation in 70% bleach (v/v) and 0.01% (v/v) Tween-20, sown in Murashige and Skoog (MS) with 1% (w/v) sucrose and either 1.2% (w/v) agar (vertical plates) or 0.7% (w/v) agar (horizontal plates). Seeds were stratified as above, and grown in 16 h light/8 h dark photoperiod at 22 °C.

### Phenotypic analysis of adult plants

Primary rosette branches (RI) and rosette leaves (RL) were counted 2 weeks after bolting of the main inflorescence. Only branches longer than 0.5 cm were quantified. To avoid variations due to flowering time (number of nodes and axillary meristems) in all experiments, branching was defined as the ratio between RI/RL. Plant height was determined 2 weeks after bolting, as the distance between the rosette and the apex of the main inflorescence. The position of plants within the growing trays was randomised to minimise environmental variation.

### Constructs

All plasmids were generated using Gateway technology (Invitrogen, Life Technologies). pDONR vectors carrying the CDS of *D14* and *SMXL7* were recombined with pDEST plasmids to generate binary vectors by LR Clonase reactions. For *35S Cauliflower Mosaic Virus Promoter* (*35S):D14:GFP* constructs we used the destination vector pGWB505 (Nakagawa *et al*., 2007) and for MultiSite Gateway Technology constructs we used pB7M34GW (Karimi *et al*., 2005). pAB118 and pAB119 were used for *LexA:CDS:mCherry* and *LexA:CDS:mCherry:GFP* constructs (Bleckmann *et al*., 2010). The CDS of *D14^H247A^* and *D14^G158E^*mutants (Yao *et al*., 2016) were provided by Dr. Ruifeng Yao. Primers used are listed in Supplemental Table 1.

### Generation of transgenic lines

Binary vectors were transformed into the *Agrobacterium tumefaciens* strain AGL-0. Transgenic Arabidopsis plants were generated by agroinfiltration using the floral dip method (Clough and Bent, 1998).

### LUC activity assays

LUC assays were performed as described (Sánchez *et al*., 2018; Sánchez *et al*., 2021). Briefly, 6 days post germination (dpg), *LUC*-expressing seedlings were placed facing up in MS-containing 96-wells plates. D-luciferin substrate (Sigma) was added to each well to a final concentration of 10 μM, and plates were pre-incubated for 2-3 h before treatment. LUC activity (counts per second, cps) was measured using a LB 960 microplate luminometer centre system (Berthold Technologies) with MikronWin 2000 software under controlled temperature (22 °C). LUC activity was measured (2 s counting time) every 10-15 min for 16 h. The LUC activity variation over time was calculated as a percentage of the t=0 signal for each plant. Then, we obtained the mean LUC activity of each treatment/genotype and represented the Relative LUC units of each treatment/genotype normalised to the mean of its mock treatment values at each time point. Experiments were performed at least twice (for n>6) or three times (for n<6).

### Time-lapse fluorescence microphotography

D14:GFP degradation assays were performed as described (Li *et al*., 2022) with 4-dpg *GFP*-expressing seedlings. Plants were treated with 5 μM GR24^5DS^ (StrigoLab) and equivalent volumes of acetone were added for mock controls. When proteasome inhibitors were used, seedlings were pre-incubated with 50 μM MG132 (PeptaNova) and 20 nM epoxomicin for 1 h before SL addition, and fresh inhibitor was added together with the hormone. Images were captured every 15-20 min for 16 h with a Microfluor Leica DMI6000B fluorescence microscope using a 10X objective and 470 nm light. Videos were obtained by Z projection with the Maximum Intensity method using the Leica Application Suite Advanced Fluorescence software (LAS-AF). GFP signal was quantified with Fiji (Schindelin *et al*., 2012) using Region of Interest (ROI) Multi Measure plugging after determining a threshold range to eliminate the background. The GFP signal variation over time was calculated as in the LUC activity assays. Each experiment was performed at least twice, with 3-4 seedlings analysed per treatment.

### Quantitative analysis of protein degradation dynamics

We defined protein half-life as the time at which plants displayed a value of LUC activity or GFP signal of 50% of that at t=0 h. When protein degradation did not reach values of 50% at the end of the assay, and therefore half-life could not be calculated, the half-life was depicted with symbols clipped at the edge of the y axis. We defined the endpoint signal as the LUC activity (%) or GFP signal (%) of each plant at the end of the assay (t=16 h for D14:GFP; t=15 h for D14:LUC; t=3 h for SMXL7:LUC) normalised to the mean of the mock treatment values.

### D14 degradation assays by immunoblot

Seven-dpg seedlings grown horizontally *in vitro* were transferred to multiwell plates with liquid MS 1% (w/v) sucrose supplemented with 5 μM GR24^5DS^, or an equivalent volume of acetone as mock control, and incubated for 8 h at 22 °C. When necessary, plants were pre-treated for 1 h with the proteasome inhibitors 50 μM MG132 (PeptaNova) and 20 nM epoxomicin. Fresh inhibitor was added together with GR24^5DS^. After treatment, seedlings were collected and frozen in liquid N_2_ for protein extraction and immunodetection. Protein was extracted in 50 mM Tris-HCl pH 7.5, 150 mM NaCl, 0.1% NP-40, 1 mM PMSF and protease inhibitor mix (Roche). Protein extracts were denatured in 5X loading buffer (1 M Tris-HCl pH 6.8, 25% (v/v) glycerol, 8% (w/v) SDS, 200 μg/ml Bromophenol Blue and 10% (v/v) β-mercaptoethanol). Samples were separated by SDS-PAGE. Proteins were transferred to a polyvinylidene difluoride (PVDF) membrane (Millipore) and probed with α-GFP-HRP antibody (1:1000, Milteny Biotec), α-ubiquitin (1:500, Enzo Life Sciences) or α-actin (1:1000, Abcam). Signal was detected using the ECL Prime Western Blotting Detection Reagent (Amersham).

### Affinity purification of ubiquitinated proteins

Seven-dpg seedlings grown horizontally in MS medium were pre-treated with the proteasome inhibitors 50 μM MG132 and 20 nM epoxomicin for 1 h and then with 10 μM GR24^rac^ (Strigolab) for further 3 h at 22°C. Seedlings were then frozen in liquid N_2_ and proteins were extracted using buffer BI (50 mM Tris-HCl pH 7.5, 20 mM NaCl, 0.1% Nonidet P-40, 5 mM ATP) plus the Protease inhibitor cocktail (Roche), 1 mM PMSF, 50 μM MG132, 10 nM Ub aldehyde (Enzo Life Sciences), 10 mM N-ethylmaleimide and 5 μM GR24^rac^. Protein extracts were incubated with 20 μL of prewashed p62 agarose (Enzo Life Sciences) or amylose resin (negative control; New England Biolabs) at 4°C for 3 h. The beads were washed twice with BI buffer and once with BI buffer supplemented with 200 mM NaCl. Proteins were eluted by boiling in 5X loading buffer and separated by SDS-PAGE. D14:GFP and total ubiquitinated proteins were detected by immunoblotting with α-GFP-HRP or α -ubiquitin respectively. Signal was detected using the ECL Prime Western Blotting Detection Reagent (Amersham) or SuperSignalTM West Femto Maximum Sensitivity Substrate (Thermo Fisher).

### Förster Resonance Energy Transfer-acceptor photobleaching (FRET-APB) assay

Leaves of 3-4-week-old *Nicotiana benthamiana* plants were agroinfiltrated and sprayed with 10 μM estradiol (Sigma) 24 h after infiltration to induce protein expression. FRET-APB assays were performed 24 h after estradiol induction on a Leica TCS SP5 laser scanning confocal microscope with a 63x/1.2NA water immersion objective as described (Nicolas *et al*., 2015). The FRET-APB wizard of LAS-AF was used with the following parameters: acquisition speed 700 Hz; pinhole 60.7 µm; image format 512×512 pixels; zoom 6X. ROIs of 6×3.5 µm were photobleached with 10 repeated exposures (laser 561 nm, 100% power level). Images were processed using LAS-AF Software. FRET Efficiency (E_FRET_%) was measured as the increase of donor (GFP) fluorescence intensity after photobleaching of the acceptor (mCherry).

### Differential Scanning Fluorimetry

Protein production in *E. coli* and DSF assays were performed as previously described (Bürger *et al*., 2019).

### Statistical analyses and bioinformatics methods

R version 3.5.3 (http://www.R-project.org/; The R Development Core Team, Preprint) and the Rcmdr package version 2.5.2 (Fox, 2005) were used for the statistical analyses. Comparisons between control and treatments were analysed by one-way ANOVA and Tukey’s test with Welch’s correction for unequal variances. Different letters denote statistical differences (*p* < 0.05). GraphPad Prism Software v. 8.3 for Windows (San Diego, California USA, www.graphpad.com) was used for data representation. The PyMOL Molecular Graphics System (Schrödinger, LLC.; https://pymol.org/) was used for molecular visualisations of 3D protein structures.

## Results

### Quantitative assays to study D14 degradation dynamics

Studies in both Arabidopsis and rice have shown that D14 is destabilised in the presence of SLs (Chevalier *et al*., 2014; Hu *et al*., 2017; Tal *et al*., 2022). To study more in detail the dynamics of D14 degradation, we generated *35S:D14:GFP* and *UB:D14:LUC* transgenic lines that allowed us to perform quantitative assays *in vivo*. Quantification of the GFP signal using time-lapse fluorescence quantitative microphotography, and LUC activity using luminescence assays could be correlated with D14 levels at different time points. We selected *35S:D14:GFP* and *UB:D14:LUC* lines that completely rescued the increased branching phenotypes of knock-out *d14-1* mutants (Supplemental Fig. 1A, B, K) and almost completely rescued the phenotype of reduced height (Supplemental Fig. 1F, G, K).

Next, we performed time-lapse fluorescence and luminescence assays to analyse, quantitatively and *in planta,* the SL-induced destabilisation of D14:GFP and D14:LUC, respectively (Fig. 1, Supplemental Fig. 2, 3, Supplemental files S1, S2). We used as reference values the protein half-life (Fig. 1B, F) and endpoint LUC or GFP signal (Fig. 1C, G). The protein half-life of D14:GFP and D14:LUC was of 3-8 h in plants treated with 5 μM GR24^5DS^ (Fig. 1A, B, D, F). However, we observed that the degradation dynamics of D14:GFP in roots and hypocotyls (which could be measured separately in the *35S:D14:GFP* lines) was not identical: D14:GFP was degraded faster (shorter half-life) and more completely (lower end-point signal) in hypocotyls than in roots (Fig. 1A-C).

**Fig. 1.**
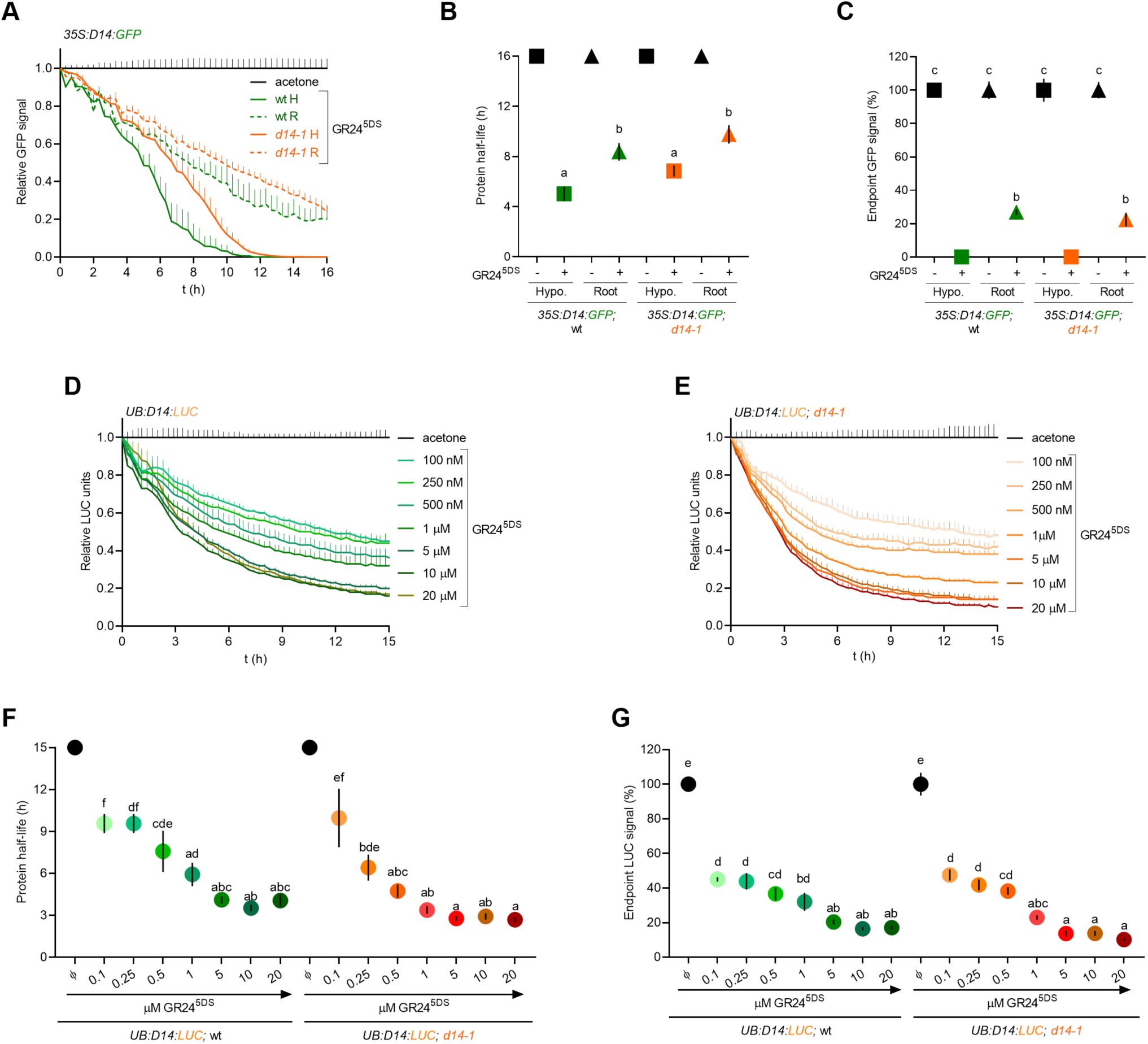
D14:GFP and D14:LUC degradation in the presence of SL. (A) Time-lapse fluorescence study of D14:GFP signal decay (relative to mock treatment values) of hypocotyls (H) and roots (R) of *35S:D14:GFP* seedlings in wild type (wt) and *d14-1* mutants treated with 5 μM GR24^5DS^ and/or the proteasome inhibitors (Inh.) 50 μM MG132 and 20 nM epoxomicin. Solvents (acetone for GR24^5DS^, DMSO for the inhibitors) were used as mock treatments (n=4). (B-C) D14:GFP protein half-life (B) and endpoint signal at 16 h (C) calculated from data in A. Square symbols represent hypocotyls, while triangles stand for roots. (D-E) LUC assays of D14:LUC signal decay (relative to mock treatment values) of *UB:D14:LUC* (D) and *UB:D14:LUC;d14-1* (E) seedlings in response to various concentrations of GR24^5DS^ or acetone as a mock treatment (n=6). (F-G) D14:LUC protein half-life (F) and endpoint signal at 15 h (G) calculated from data in D and E. Data shown as mean ± SE. Symbols clipped at the edge of the Y axis indicate conditions in which protein half-life is longer than 16 h (B) or 15 h (F). Different letters denote statistical differences in one-way ANOVA with post hoc Tukey test, *P* < 0.05.

We assayed the D14:LUC activity with concentrations of GR24^5DS^ ranging from 100 nM to 20 μM, and found that the protein was destabilised in a dose-dependent manner between 100 nM and 1-5 μM GR24^5DS^, concentrations at which the shortest half-life and most complete degradation (lowest endpoint LUC signal) were observed. Higher hormone concentrations did not lead to faster or more complete D14:LUC degradation (Fig. 1D-G). In general, the dynamics of degradation were not significantly different in the wild type and in *d14-1* mutant backgrounds either for D14:GFP (Fig. 1A-C) or for D14:LUC (Fig. 1D-G).

### Degradation of D14:GFP is only partly mediated by the 26S proteasome

Previous studies have suggested that the proteasome is the pathway for D14 destabilisation. (Chevalier *et al*., 2014; Hu *et al*., 2017). Proteins targeted for proteasomal degradation are polyubiquitinated for recognition by the proteasome machinery. To test whether D14:GFP was ubiquitinated upon SL treatment in Arabidopsis, we treated 7-dpg *35S:D14:GFP* seedlings with GR24, affinity-purified ubiquitinated proteins from their extracts using a Ubiquitin (Ub)-binding resin, and performed α-GFP immunoblots. We detected D14:GFP and higher molecular weight bands corresponding to different D14:GFP poly-ubiquitinated forms (Ub(n):D14:GFP; Fig. 2A). This result confirmed that, like rice D14, Arabidopsis D14:GFP is polyubiquitinated in the presence of exogenous SLs in Arabidopsis.

**Fig. 2.**
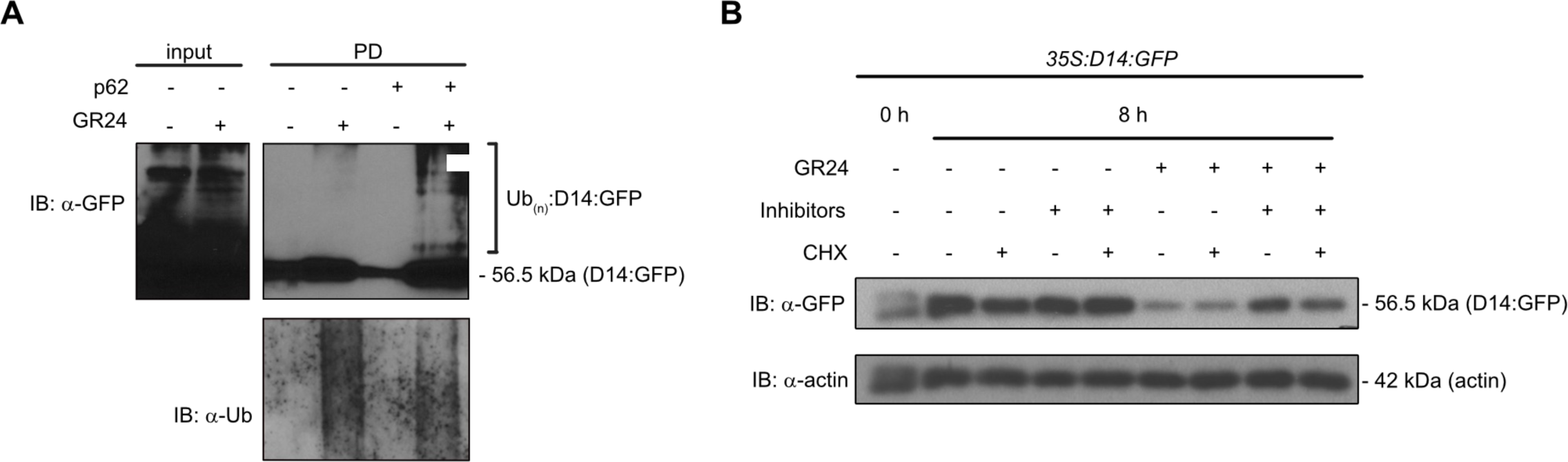
SL-induced D14:GFP degradation is only partly mediated by the 26S proteasome. (A) D14:GFP is poly-ubiquitinated in response to GR24. Seven dpg, *35S:D14:GFP* plants were incubated with 10 μM GR24 and 50 μM MG132 for 3 h. Poly-ubiquitinated proteins were affinity purified from protein extracts by incubation with Ub-binding p62 resin (+) or empty agarose resin (negative control; −). Immunoblots were performed with α-ubiquitin antibodies to detect total ubiquitinated proteins or α-GFP to detect D14:GFP and its ubiquitinated forms (Ub(n):D14:GFP). The molecular mass of the D14:GFP protein is indicated. (B). Immunoblot of 7 dpg *35S:D14:GFP* seedlings treated for 8 h with 5 μM GR24 and the proteasome inhibitors (Inhibitors) 50 μM MG132 and 20 nM epoxomicin. In the indicated lanes, 50 μM cycloheximide (CHX) was added to prevent *de novo* protein synthesis. Solvents (acetone for GR24, DMSO for CHX and Inhibitors) were used as mock treatments. An α-actin antibody was used for loading control. The molecular masses of D14:GFP and actin are indicated.

It has been reported that SL-induced D14 degradation can be suppressed by treatments with the 26S proteasome inhibitor MG132 (Chevalier *et al*., 2014; Hu *et al*., 2017). To confirm this, we performed immunoblots with 7-dpg *35S:D14:GFP* seedlings treated with GR24^5DS^ both with and without proteasome inhibitors (MG132 and epoxomicin). Plants treated with GR24^5DS^ and proteasome inhibitors displayed higher levels of D14:GFP than those without inhibitors, but some degradation of D14:GFP was still evident when compared to plants not treated with GR24^5DS^ (Fig. 2B). Moreover, time-lapse fluorescence microphotography seemed to indicate that the D14:GFP degradation dynamics are delayed, but not abolished by the inhibitors (Fig. 3A, B), and that the half-life (Fig. 3F) and endpoint signal (Fig. 3G) parameters did not always capture significant differences between treated and untreated plants. These results point to the existence of a proteasome-independent mechanism contributing to D14:GFP degradation along with the proteasome-dependent pathway.

**Fig 3.**
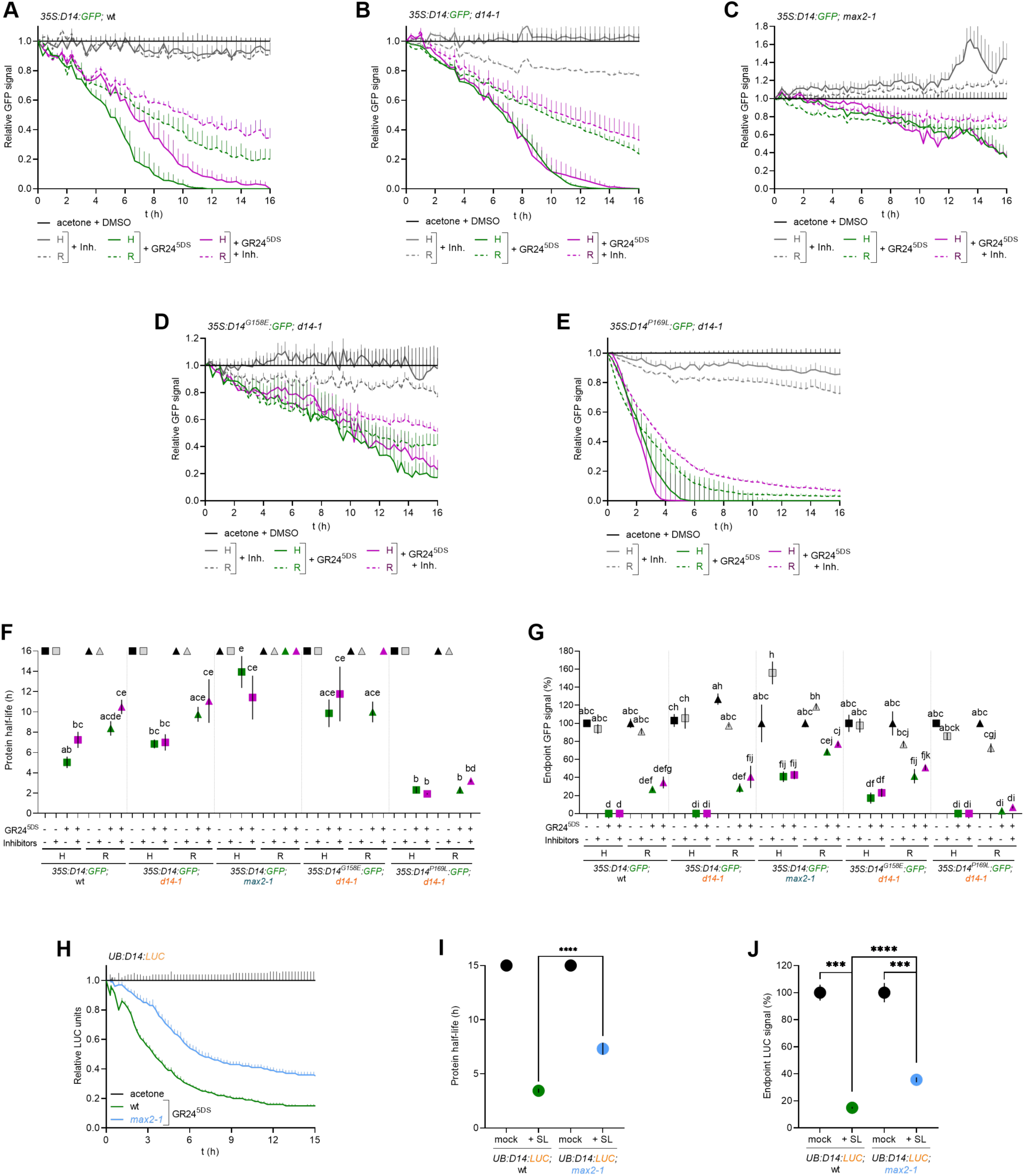
MAX2-independent D14 degradation. (A-D) Time-lapse fluorescence study of signal decay (relative to mock treatment values) of hypocotyls (H) and roots (R) treated with 5 μM GR24^5DS^ and/or the proteasome inhibitors (Inh.) 50 μM MG132 and 20 nM epoxomicin for D14:GFP (A), D14:GFP (*d14-1* background) (B), D14:GFP (*max2-1* background) (C), D14^G158E^:GFP (*d14-1* background) (D) and D14^P169L^:GFP (*d14-1* background) (E). Solvents (acetone for GR24^5DS^, DMSO for the Inhibitors) were used as mock treatments (n=3). (E-F) D14:GFP, D14^G158E^:GFP and D14^P169L^:GFP protein half-life (F) and endpoint signal at 16 h (G) calculated from data in A-D. Square symbols represent hypocotyls, while triangles stand for roots. (H) LUC assay of D14:LUC signal decay (relative to mock treatment values) of *UB:D14:LUC* seedlings in wild type (wt) and *max2-1* mutants treated with 5 μM GR24^5DS^ or acetone as mock control (n=6-9). (I-J) D14:LUC half-life (I) and endpoint signal at 15 h (J) calculated from data in H. Data shown as mean ± SE. Symbols clipped at the edge of the Y axis (F, G) indicate conditions in which protein half-life is longer than 16 h or 15 h. Different letters denote statistical differences in one-way ANOVA with post hoc Tukey test, *P* < 0.05; asterisks indicate significant differences between pairs in Student’s t test (**** *P* < 0.0001; *** *P* < 0.001).

### The strigolactone signalling complex SCF^MAX2^ contributes to, but is not essential for D14 degradation

Previous studies have shown that SL-induced D14 degradation is largely compromised in *max2* mutants (Chevalier *et al*., 2014; Hu *et al*., 2017), suggesting that the SCF^MAX2^ complex is pivotal in this process. Therefore, we quantitatively investigated the role of *MAX2* in more detail, by performing D14:GFP stability assays in *35S:D14:GFP;max2-1* lines. Remarkably, we found that, in *max2* mutants, there was still a certain level of D14:GFP degradation induced by GR24^5DS^, although reduced as compared to the degradation observed in the wild type. This degradation was largely unaffected by proteasomal inhibitors (Fig. 3C, F, G). We further confirmed these results by LUC assays performed in *UB:D14:LUC;max2-1* (Fig. 3H-J).

As a complementary approach, we analysed, in SL-treated plants, the stability of two D14 mutant proteins whose capability to interact with MAX2 is compromised due to amino acid substitutions at the lid domain, a region involved in the D14-MAX2 interaction. One of these proteins was D14^G158E^ (Supplemental Fig. 4A, B), unable to interact with the MAX2 rice ortholog D3 in pull-down assays (Yao *et al*., 2016). In *35S:D14^G158E^:GFP;d14-1* transgenic lines treated with GR24^5DS^, we detected a significant although not complete degradation of D14^G158E^:GFP, similar to that observed in *max2* mutants, and also insensitive to proteasome inhibitors (Fig. 3D, F, G). DSF assays with increasing concentrations of GR24 indicated that D14^G158E^ gets destabilised only at much higher concentrations of GR24^5DS^ than wild-type D14 (Supplemental Fig. 5A, B), which may explain the suboptimal degradation of the protein.

Next we analysed the SL-induced proteolysis of D14^P169L^:GFP, another mutant protein with an amino acid substitution colocalizing with the area of D14-MAX2 interaction on the external surface of the lid domain (Chevalier et al., 2014; Yao et al., 2016; Supplemental Fig. 4C, D). In *35S:D14^P169L^:GFP;d14-1* transgenic lines, we detected a fast and significant protein degradation in response to GR24^5DS^, insensitive to proteasome inhibitors (Fig. 3E-G; Li *et al*., 2022). These degradation patterns were indistinguishable in wild-type and in *max2* mutant backgrounds (*35S:D14^P169L^:GFP;max2-1*) (Supplemental Fig. 6A-C), consistently with a lack of interaction of D14^P169L^ with MAX2.

All these results suggest that SLs can still promote the degradation of D14 in the absence of an interaction with MAX2. This degradation may occur through a mechanism independent of the proteasome.

### The interaction of D14 with the SL signalling repressors SMXL6, SMXL7 and SMXL8 contributes to, but is not essential for D14 degradation

In the presence of SLs, the SMXLs are rapidly recruited to the SL-D14-SCF^MAX2^ complex and targeted for proteasomal degradation. We assessed whether this was a prerequisite for D14 destabilisation by performing D14:LUC degradation assays in the triple mutant *smxl6-4 smxl7-3 smxl8-1* (*s678)* (*UB:D14:LUC;s678* lines). D14:LUC was destabilised in the *s678* mutant background with degradation dynamics almost identical to those observed in wild type (Fig. 4A-C). This indicates that these SMXLs are not essential for SL-induced D14 degradation. To assess whether the SMXLs-independent D14 degradation occurred via SCF^MAX2^, we studied D14:LUC degradation in the quadruple mutant *s678;max2-1* (*UB:D14:LUC;s678;max2*). In *s678;max2*, D14:LUC half-life was longer than in wild type or *s678* but shorter than in *max2-1* (Fig. 4A-C). This may indicate that, in the absence of the SMXLs, D14 degradation can still occur both *via* the faster SCF^MAX2^ pathway (*s678*) or via a slower MAX2-independent pathway (*s678;max2*). Furthermore, the observation that in *max2* mutants D14:LUC half-life is longer than in *s678;max2* suggests that the interaction of the SMXLs with SL-D14 may interfere with the MAX2-independent pathway of D14 degradation, perhaps by making D14 less accessible to the alternative proteolytic machinery.

**Fig. 4.**
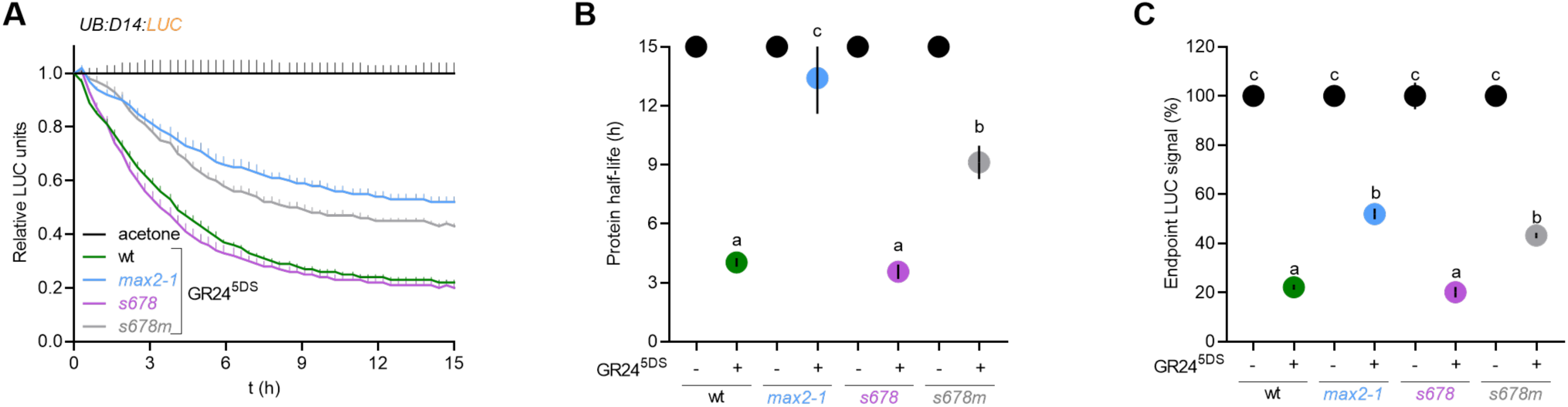
D14:LUC degradation does not require the SMXLs. (A) LUC assay of D14:LUC signal decay (relative to mock treatment values) of *UB:D14:LUC* in wild type (wt), *max2-1*, *smxl6/7/8* triple mutant (*s678*) and *s678max2-1* (*s678m*) quadruple mutant seedlings treated with 5 μM GR24^5DS^ or acetone as a mock control. (B-C) D14:LUC protein half-life (B) and endpoint signal at 15 h (C) calculated from data in A. Data shown as mean ± SE (n=12). Symbols clipped at the edge of the Y axis indicate conditions in which protein half-life is longer than 15 h. Different letters denote statistical differences in one-way ANOVA with post hoc Tukey test, *P* < 0.05.

### Relationship between D14 and SMXL7 degradation patterns and SL signalling status

Next, we investigated whether the extent of D14 degradation was strongly associated with SMXL7 degradation and SL signalling in Arabidopsis. For this we used *UB:SMXL7:LUC* transgenic lines that we combined with *d14-1* and *max2-1* mutants and with *D14:GFP*-expressing transgenic lines.

First, we confirmed that SMXL7:LUC degradation was strictly dependent on D14, the only reported SL receptor in Arabidopsis. Indeed, *d14-1* mutants displayed no degradation of SMXL7:LUC upon GR24^5DS^ treatment (Fig. 5A-C). This contrasted with the efficient destabilisation of D14:LUC in *s678* mutants (see above, Fig. 4A).

**Fig. 5.**
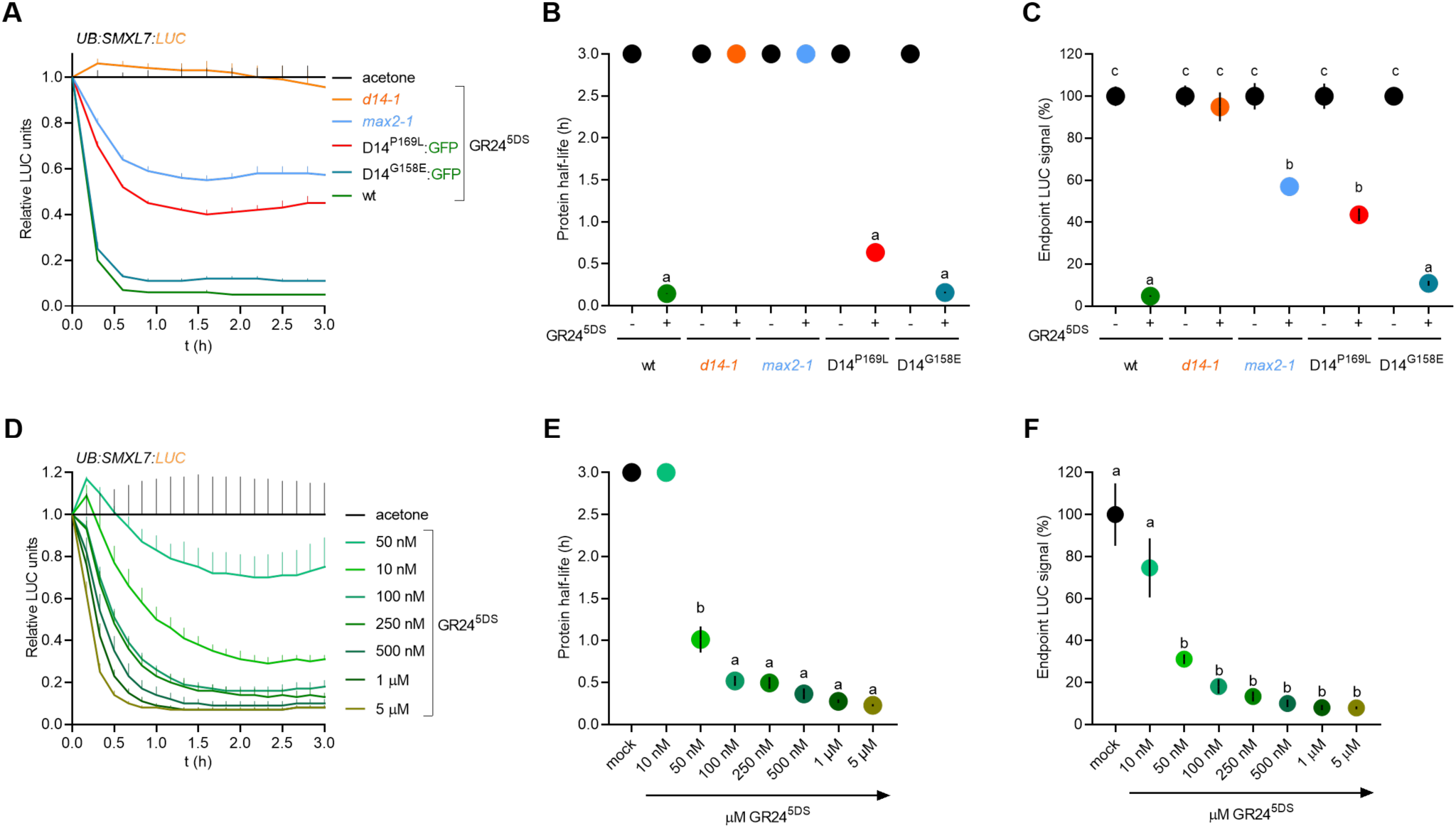
Relationship between SL signalling and D14 degradation. (A) LUC assay of SMXL7:LUC signal decay (relative to mock treatment values) of *UB:SMXL7:LUC* wild type (wt), *d14-1, max2-1, 35S:D14^P169L^:GFP;d14-1* (D14^P169L^) or *35S:D14^G158E^:GFP;d14-1* (D14^G158E^) seedlings treated with 5 μM GR24^5DS^ or acetone as mock (n=6). SMXL7:LUC protein half-life (B) and endpoint signal (C) at 3 h calculated from data in A. Data shown as mean ± SE. (D) LUC assay of SMXL7:LUC signal decay (relative to mock treatment values) of *UB:SMXL7:LUC* seedlings in response to various concentrations of GR24^5DS^ or acetone as a mock control (n=4). SMXL7:LUC protein half-life (E) and endpoint signal (F) at 3 h calculated from data in D. In B and E, symbols clipped at the edge of the Y axis indicate conditions in which protein half-life is longer than 3 h. Different letters denote statistical differences in one-way ANOVA with post hoc Tukey test, *P* < 0.05.

Next, we studied whether there was a correlation between the SL-induced degradation of D14:GFP/LUC and that of SMXL7:LUC in several conditions and genetic backgrounds analysed in this work. In general, SMXL7 degradation efficiency paralleled that of D14. SMXL7:LUC was destabilised by GR24^5DS^ in a dose-dependent manner (Fig. 5D-F) although at much lower SL concentrations than D14 (Fig. 1D, E). Also, the partial degradation of D14 in *max2-1* (Fig. 3C, H) was mirrored by a partial degradation of SMXL7:LUC (*UB:SMXL7:LUC;max2-1* lines) (Fig. 5A-C) supporting the possibility that SMXL7:LUC can also be targeted from degradation upon interaction with SL-D14 in the absence of MAX2.

However, in certain scenarios, the efficiency of SL-induced degradation of D14 and SMXL7 were largely different. For instance, we showed a noticeable but very reduced degradation of D14^G158E^:GFP as compared to D14:GFP (Fig. 3D, F, G). We combined *UB:SMXL7:LUC;d14-1* with *35S:D14^G158E^:GFP;d14-1* and studied SMXL7:LUC degradation in the F1 (*35S:D14^G158E^:GFP/+;UB:SMXL7:LUC/+;d14-1*). These plants were in a *d14-1* mutant background, in which SMXL7:LUC could only interact with D14^G158E^:GFP. Remarkably, SMXL7:LUC displayed a fast and efficient degradation indistinguishable from that in the wild type, in which SMXL7:LUC interacts with a wild-type D14 (Fig. 5A-C). Conversely, although D14^P196L^:GFP degradation was slightly faster than that of D14:GFP (Figure 3E-G), the SMXL7:LUC degradation in *35S:D14^P169L^:GFP/+; UB:SMXL7:LUC/+*; *d14-1* was significantly less efficient than in the wild-type background (Fig. 5A-C). These results indicate that the degradation of the repressor of the SL pathway SMXL7 and that of the receptor D14 are not necessarily correlated in a quantitative manner.

Moreover, D14 degradation efficiency was not consistently aligned with the status of SL signalling, as measured by the phenotype of the lines studied. For example, despite the inefficient degradation of D14^G158E^:GFP, the *35S:D14^G158E^:GFP*;*d14-1* lines had an almost wild-type branching and height phenotypes (Supplemental Fig. 1D, I, K) in agreement with the wild-type SMXL7:LUC degradation patterns (Fig. 5D-F). In contrast, although D14^P196L^:GFP degradation is slightly faster and more efficient than that of D14, the *35S:D14^169E^:GFP;d14-1* plants have a phenotype similar to *d14-1* of increased branching and reduced height (Supplemental Fig. 1E, J, K) and a partial degradation of SMXL7 (Fig. 5A-C). The latter observation also shows that SMXL7 degradation is in line with SL signalling and that an incomplete degradation of SMXL7 is insufficient to achieve full SL signalling, at least in terms of the control of branching patterns and plant height. However, D14 degradation does not provide information about, nor is necessarily associated with the success of SL signalling.

### SL binding but not hydrolysis is required for D14 degradation

D14 is an unusual hormone receptor that not only binds but also hydrolyses the bound SL molecule. This has raised the question of whether binding is sufficient, or hydrolysis is also required for SL signalling (Yao *et al*., 2016; Saint Germain *et al*., 2016; Seto *et al*., 2019). Likewise, it is yet unclear whether both binding and hydrolysis are needed for SL-induced D14 degradation. Rice transgenic lines expressing *D14* genes carrying mutations in the catalytic triad, which preclude D14 from hydrolysing the SL ligand, showed a drastic decrease in D14 degradation (Hu *et al*., 2017). This led to the proposal that hydrolysis was necessary for SL-mediated destabilisation of D14.

Therefore, we assayed the stability of the D14^H247A^ mutant that bears a histidine for alanine substitution at the catalytic triad and has been suggested to be incapable to hydrolyse SLs (Yao *et al*., 2016). Time-lapse studies of *35S:D14^H247A^:GFP;d14-1* transgenic lines showed that D14^H247A^:GFP was fully resistant to GR24^5DS^-induced degradation *in planta* (Supplemental Fig. 6D, E). Moreover, the *35S:D14^H247A^:GFP;d14-1* lines had a strong *d14* mutant phenotype (Supplemental Fig. 1C, H, K). However, more recently it has been shown that SL binding is also impaired in D14^H247A^ (Seto *et al.,* 2019). We confirmed this observation by DSF assays with GR24^5DS^ and D14^H247A^:GFP, which showed no thermal shift (Supplemental Fig. 5A, C). These results indicate that mutations at the catalytic triad of D14 can affect not only hydrolysis but also hormone binding, and underscore the need for caution when interpreting the behaviour of mutants in the catalytic triad of D14. In any case, this also confirms that SL binding is essential for D14 degradation.

Therefore, instead of catalytic triad mutants, we used an alternative, pharmacological approach to investigate the requirement for SL hydrolysis: a non-hydrolysable SL derivative, carba-GR24 (Thuring *et al.,* 1997; Prandi & McErlean, 2019). We performed LUC assays in *UB:D14:LUC* lines treated with carba-GR24 or GR24, and found that D14:LUC was still degraded in the presence of carba-GR24 (Fig. 6A, C) although at a slower rate and using higher concentrations of the compound (20 μM) than when treated with GR24 (Fig. 6A, B). These results indicate that hormone hydrolysis is not essential for D14 degradation, although it reduces protein half-life during the process. Moreover, 20 μM carba-GR24 also promoted SMXL7:LUC degradation although, at significantly slower rate and with lower efficiency (up to 45% of its original values) than GR24 (Fig. 6D-F). Consistently, FRET assays showed that carba-GR24 enabled the interaction of D14 with SMXL7 (Fig. 6G). These results suggest that D14 and SMXL7 degradation (and probably SL signalling) can also occur without SL hydrolysis, although in a less efficient manner.

**Fig. 6.**
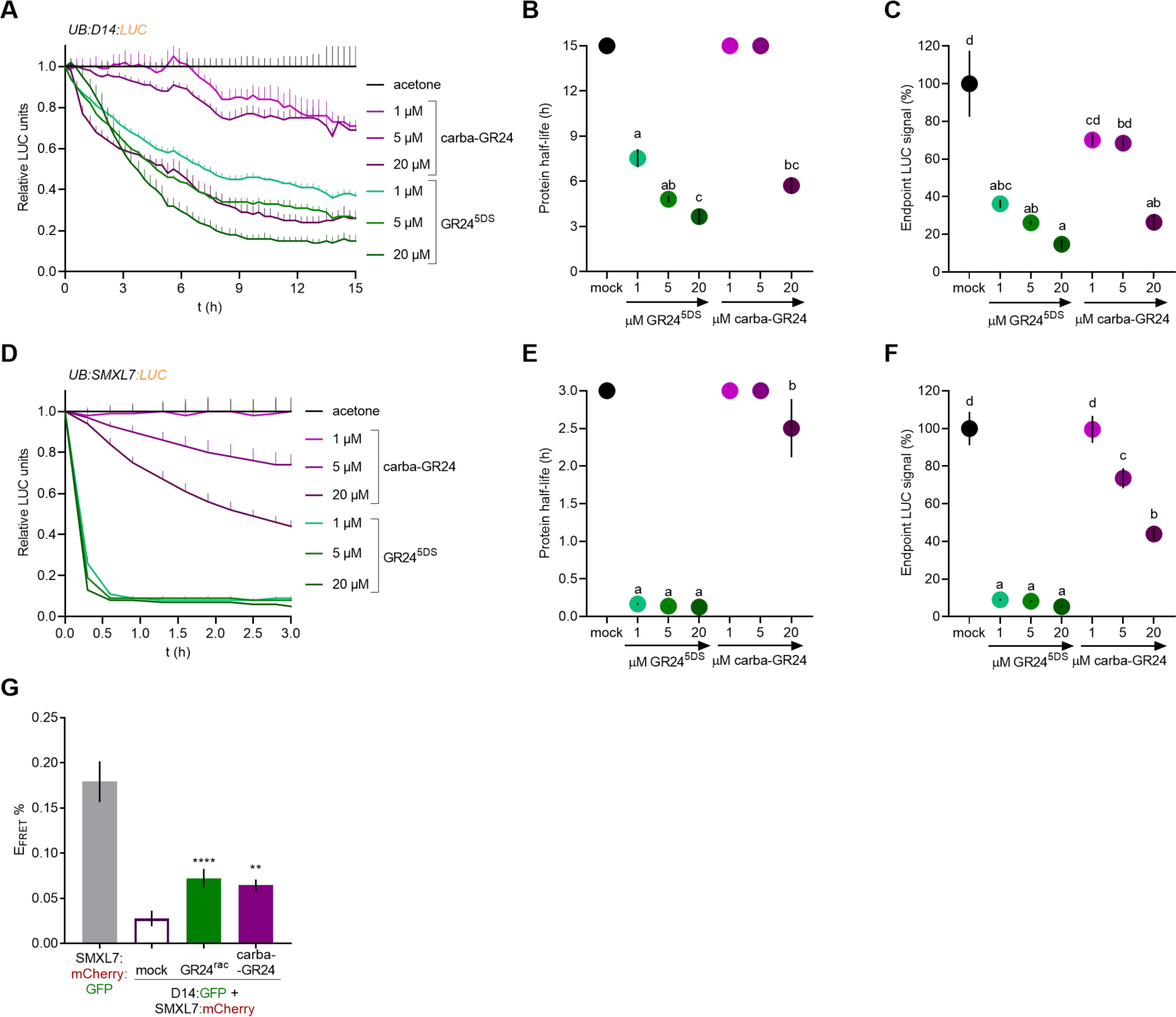
SL binding but not hydrolysis is necessary for D14 degradation. (A) LUC assays of D14:LUC signal decay (relative to mock treatment values) in *UB:D14:LUC* seedlings treated with various concentrations of GR24^5DS^, the non-hydrolysable SL carba-GR24, or acetone (mock). (B-C) D14:LUC protein half-life (B) and endpoint signal at 15 h (C) calculated from data in A (n=6). Symbols clipped at the edge of the Y axis indicate conditions in which protein half-life is longer than 15 h. Different letters denote statistical differences in one-way ANOVA with Welch’s test. (D) LUC assay of SMXL7:LUC signal decay (relative to mock treatment values) of *UB:SMXL7:LUC* seedlings in response to various concentrations of GR24^5DS^, the non-hydrolysable SL C.L. or acetone as mock (n=6). (E-F) SMXL7:LUC protein half-life (E) and endpoint signal at 3 h (F) calculated from data in D. (D) FRET-APB of SMXL7:mCherry and D14:GFP after 30 min of treatment with 20 µM GR24^rac^ or 50 µM C.L. E_FRET_% is calculated as the relative increase in GFP fluorescence intensity after photobleaching of the mCherry acceptor (n=6-12 nuclei). Positive control, intramolecular FRET of a SMXL7:mCherry:GFP protein. Data shown as mean ± SE. In B and E, symbols clipped at the edge of the Y axis indicate conditions in which protein half-life is longer than 15 h or 3 h. Different letters denote statistical differences in one-way ANOVA with post hoc Tukey test, *P* < 0.05; asterisks indicate significant differences between pairs in Student’s t test (**** p<0.0001; ** p<0.01).

## Discussion

Extensive work performed in Arabidopsis and rice allowed the generation of a working model of the molecular mechanisms controlling SL perception and signalling. Compelling evidence indicates that D14 is the only receptor for the endogenous SLs, and that it also hydrolyses the hormone. Moreover, the pivotal role of the SCF^MAX2^ complex in the ubiquitination and proteasomal degradation of the transcriptional repressors SMXL6, SMXL7 and SMXL8 has been largely proven. The observation that D14 is itself destabilised in the presence of SLs has raised the question of whether a potential feedback regulatory mechanism of SL perception is controlled by the same machinery that causes SMXLs degradation. Previous work has suggested that this is the case (Chevalier *et al.,* 2014; Hu *et al.,* 2017), at least to a significant extent. Here we present evidence of a more complex scenario in which, in addition to the SCF^MAX2^-dependent pathway of D14 degradation via the proteasome in Arabidopsis, alternative pathways may contribute to limiting D14 protein levels in the presence of SLs.

Indeed, SCF^MAX2^ is not strictly required for SL-induced D14 degradation in Arabidopsis: in *max2* mutants D14 gets destabilised, although less efficiently than in wild type. Moreover, D14 mutant proteins predicted to be incapable of interacting with MAX2, namely D14^G158E^ and D14^P169L^ (Yao *et al.,* 2016; Chevalier *et al.,* 2014), still display a significant degradation in response to SLs. This indicates that alternative pathways exist, although not as effective as that of SCF^MAX2^. Other E3 ligases could promote D14 degradation. For instance, a physical interaction has been reported between rice D14 and the RING-finger ubiquitin E3 ligase SDEL1 under phosphate (Pi) deficiency, which promotes degradation of SPX DOMAIN-CONTAINING PROTEIN 4 and release of PHOSPHATE STARVATION RESPONSE PROTEIN 2, thus improving Pi acquisition and translocation (Gu *et al*., 2023). It remains to be tested whether D14 undergoes ubiquitination and degradation in this system. Nevertheless, the MAX2-independent pathway of D14 degradation observed in this work may not involve the proteasome, as the destabilisation of D14:GFP in *max2*, and that of D14^G158E^ and D14^P169L^ are not blocked by proteasome inhibitors. Alternative proteolytic pathways need to be explored to fully understand this phenomenon.

In addition, it has been proposed that the degradation of rice D14 is tightly coupled to the degradation of D53, the rice ortholog of the SMXLs. This was based on the observation that D14:GFP destabilisation is significantly impaired (although not abolished) in the SL-insensitive *d53* mutants (Hu *et al*., 2017). In Arabidopsis, we have found that D14 degradation is similar in wild type and *smxl678* mutants, which indicates that lack of SMXLs does not influence D14 destabilisation. However, SMXLs may delay D14 degradation through the MAX2-independent pathway, as inferred by the faster degradation of D14 in the *smxl678;max2* mutants than in the *max2* mutants. Without a functional SCF^MAX2^, the interaction of the SMXLs with SL-D14 could in part protect D14 from the proteolytic machinery.

Furthermore, we have observed plants displaying, on one hand, an efficient wild type-like degradation of SMXL7 together with a poor degradation of D14^G158E^ and, on the other hand, a limited degradation of SMXL7 along with a fast and efficient degradation of D14^P169L^. This indicates that both responses are not always necessarily coupled in a quantitative manner. The D14^G158E^ and D14^P169L^ mutants may be affected differently in the SL-induced conformational changes of D14 (as DSF studies have shown for D14^G158E^). This may impact their interactions with the SMXLs and with the proteolytic machineries, thereby facilitating some responses and preventing others.

Nevertheless, whereas SMXL7 degradation is strongly associated with the status of SL signalling, D14 degradation does not seem to provide information about the success of SL signalling. This is clearly illustrated by the fast-degrading D14^P169L^ mutant protein, which is unable to rescue the *d14-1* phenotype (Chevalier *et al.,* 2014). Moreover, a partial degradation of SMXL7 (as observed in *max2* mutants and in *35S:D14^P169L^:GFP;d14-1* lines) is insufficient to achieve full SL signalling, at least in terms of branching and height phenotypes.

The requirement of SL hydrolysis for the degradation of D14 was also an open question. Hu *et al*. (2017) had shown that rice D14:GFP proteins carrying mutations in residues of the catalytic triad essential for the hydrolase activity display severely impaired D14 degradation, which led them to propose that hydrolysis is essential for this process. However, it is becoming increasingly clear that some D14 catalytic triad mutants are unable to bind SLs (Zhao *et al.,* 2015; Seto *et al.,* 2019, this work), which highlights the importance of being cautious when interpreting results regarding catalytic triad mutants. Our results demonstrate that the non-hydrolysable SL carba-GR24 promotes degradation of D14, interaction between D14 and SMXL7, and destabilisation of SMXL7. These findings strongly suggest that SL hydrolysis is not essential in this process, and are in line with the proposal that the intact SL molecule serves as the active compound that triggers conformational changes in D14 and initiates not only SL signalling (Seto *et al.,* 2019) but also D14 degradation.

Our studies with D14 mutant proteins also provide information about the functional domains of the D14 protein and its interactions with other components of the SL signalling pathway. For instance, the observation that in *35S:D14^G158E^:GFP/+;UB:SMXL7:LUC/+;d14-1* plants, SMXL7:LUC displays almost wild-type degradation dynamics suggests that the mutant protein D14^G158E^:GFP (unable to interact with D3/MAX2, Yao *et al.,* 2016), could still interact with SMXL7:LUC and target it for degradation through a yet unknown but rather efficient mechanism.

D14^P169L^, whose mutation is also located in the lid domain, has been proposed to fail to interact with MAX2 (Chevalier *et al*., 2014). Although our attempts to directly test D14 and MAX2 interactions using YTH and FRET assays were unsuccessful, the observation that D14^P169L^:GFP degradation dynamics are the same in wild type and *max2* mutants support the possibility of a lack of interaction between D14^P169L^ and MAX2. Remarkably, unlike D14^G158E^, the D14^P169L^ mutant is very inefficient in targeting SMXL7:LUC for degradation. D14^P169L^ seems to interact with other SMXL proteins, perhaps with higher affinity than with SMXL7 (Li *et al*., 2022). This might account for the incomplete degradation of SMXL7:LUC and the pronounced mutant phenotype observed in the mutant lines.

In conclusion, our investigation into the degradation patterns of wild-type and mutant D14 proteins *in planta* reveals novel regulatory features of SL-induced D14 destabilisation. Beyond the canonical pathway involving ubiquitination by SCF^MAX2^ and subsequent proteasomal degradation, our findings highlight the existence of additional mechanisms that operate independently of MAX2 and the proteasomal machinery. Remarkably, these MAX2-independent mechanisms can also effectively target SMXL7 for degradation. Moreover, SMXLs are not required for SL-induced degradation of D14. Advanced and sensitive proteomic assays and genetic analysis may help identify additional interactors of D14 involved in its destabilisation upon SL binding. These new interactors may shed light on the crosstalk between SL signalling and other signalling pathways, contributing to a more comprehensive view of the plant hormone response networks.

## Supporting information

Supplemental figures Sanchez Martin-Fontecha et al

Supplemental file S1 Sanchez Martin-Fontecha et al

Supplemental file S2 Sanchez Martin-Fontecha et al

## Acknowledgements

We thank V. Rubio and E. Iniesto for advice on protein ubiquitination assays, M. Nicolas for guidance with APB-FRET studies, R. García for technical support with LUC assays and plant phenotyping. The research was supported by grants from Spanish Ministry of Economy (MINECO) and fondos FEDER PID2020-112779RB-I00/AEI/ 10.13039/501100011033, BIO2017-84363-R, BIO2017-84066-R and BIO2014-57011-R]. Elena Sánchez Martín-Fontecha had an FPI (MINECO) fellowship BES-2015-074233, funded by MCIN/AEI/10.13039/501100011033 and by ESF Invest in Your Future.

## Author contributions

PC designed research, ESMF and MB performed research, PC, ESMF, FC, MB and CP analysed data, CP provided materials, PC, ESMF and FC wrote article.

## Figure legends

**Supplemental Table S1. Primers used in this work**

**Supplemental Fig. 1. Functional analysis of D14:GFP, D14:LUC and D14:GFP mutant proteins.** (A-J) Shoot branching (A-E) and height (F-J) phenotypes of wild type (wt), *d14-1* and *35S:D14:GFP;d14-1* lines (A, F), *UB:D14:GFP;d14-1* lines (B, G), *35S:D14*^H247A^*:GFP;d14-1* (C, H), *35S:D14^G158E^:GFP;d14-1* (D, I) and *35S:D14^P169L^:GFP;d14-1* (E, J) two weeks after bolting. Branching is denoted as the number of primary rosette branches/number of rosette leaves (RI/RL). Violin plots represent data distribution. Black dotted lines indicate median values; grey dotted lines delimit the interquartile range (n=11-16). (K) Images of representative individuals from each line. Different letters denote statistical differences in one-way ANOVA with post hoc Tukey test, *P* < 0.05. For assays with *35S:D14:GFP;d14-1,* line L18 was used.

**Supplemental Fig. 2. D14:GFP time-lapse assay in hypocotyls.**

Time series of fluorescence images of the hypocotyl of *35S:D14:GFP;d14-1* seedlings treated with 5 μM GR24^5DS^ (A) or acetone as mock treatment (B). Images were taken every 20 minutes up to 16 hours. Scale bar = 500 µm.

**Supplemental Fig. 3. D14:GFP time-lapse assay in roots.**

Time series of fluorescence images of the root of *35S:D14:GFP;d14-1* seedlings treated with 5 μM GR24^5DS^ (A) or acetone as mock treatment (B). Images were taken every 20 minutes up to 16 hours. Scale bar = 500 µm

**Supplemental Fig. 4. Localization of the G158 and P169 residues possibly involved in the D14-MAX2 interaction.** 3D molecular visualisation as a surface representation of the D14 protein in modified PDB:4IH4 (Zhao *et al*., 2013) (A,C) and PDB:5HZG (Yao *et al*., 2016) (B,D). The α/β hydrolase core domain is coloured in grey; the lid domain in salmon; residues of the catalytic triad in blue; residues G158 and P169 are highlighted in red. In B and D, the ribbon representation of the rice MAX2 orthologue OsD3 is coloured in green.

**Supplemental Fig. 6. Evaluation of D14, D14^H247A^ and D14^G158E^ protein stability in the presence of GR24^5DS^**. In the DSF assays, the melting temperature (Tm) profiles of the D14 (A), D14^G158E^ (B) and D14^H247A^ (C) proteins, each isolated from *E. coli*, were monitored under increasing concentrations of GR24^5DS^, and thermal denaturation shifts observed in these proteins were recorded.

**Supplemental Fig. 5. Analysis of D14^H247A^:GFP and D14^G158E^:GFP stability in response to GR24^5DS^.** (A) Time-lapse fluorescence study of D14^P169L^:GFP signal decay (relative to mock treatment values) of hypocotyls (H) and roots (R) of *35S:D14^P169L^:GFP;d14-1* seedlings treated with 5 μM GR24^5DS^ or acetone (mock). (B-C) D14^P169L^:GFP protein half-life (B) and endpoint signal at 16 h (C) calculated from data in A. Square symbols represent hypocotyls, while triangles stand for roots. (D) Time-lapse fluorescence study of D14^H247A^:GFP signal decay (relative to mock treatment values) of hypocotyls (H) and roots (R) of *35S:D14^H247A^:GFP;d14-1* seedlings treated with 5 μM GR24^5DS^ or acetone (mock). (E) D14^H247A^:GFP endpoint signal at 16 h calculated from data in D. Square symbols represent hypocotyls; triangles, roots. Data shown as mean ± SE (n=3). In B, symbols clipped at the edge of the Y axis indicate conditions in which protein half-life is longer than 16 h. Different letters denote statistical differences in one-way ANOVA with post hoc Tukey test, *P* < 0.05.

**Supplemental File S1. D14:GFP time-lapse assay in hypocotyls.**

**Supplemental File S2. D14:GFP time-lapse assays in roots.**

