## Supplemental figures Sanchez Martin-Fontecha et al for "Novel Mechanisms of Strigolactone-Induced DWARF14 Degradation in *Arabidopsis thaliana*"

| Name | Sequence | Use |
| --- | --- | --- |
| attB1-D14 | ggggacaagtttgtacaaaaaagcaggctTAATGAGTCAACACAACATCT | Cloning of D14$^{H247A}$ and D14$^{G158E}$ |
| attB2-D14ΔSTOP | ggggaccactttgtacaagaaagctgggtCGGCCCGAGGAAGAGCTCGCC | Cloning of D14$^{H247A}$ and D14$^{G158E}$ |
| max2F | CGAGAATTTTTGACGCCATT | Genotyping of *max2-1* |
| max2R | GTGGTGGCCAATAATCAAGC | Genotyping of *max2-1* |
| d14-A | GTACTGACACATTCATTGGAACCAC | Genotyping of *d14-1* |
| d14-B | AGCCGGCACAGAAACATCCTTCGC | Genotyping of *d14-1* |
| LB-1 | GGCAATCAGCTGTTGCCCGTCTCACTGGTG | Genotyping of *d14-1* |

**Supplemental Table 1. Primers used in this work**

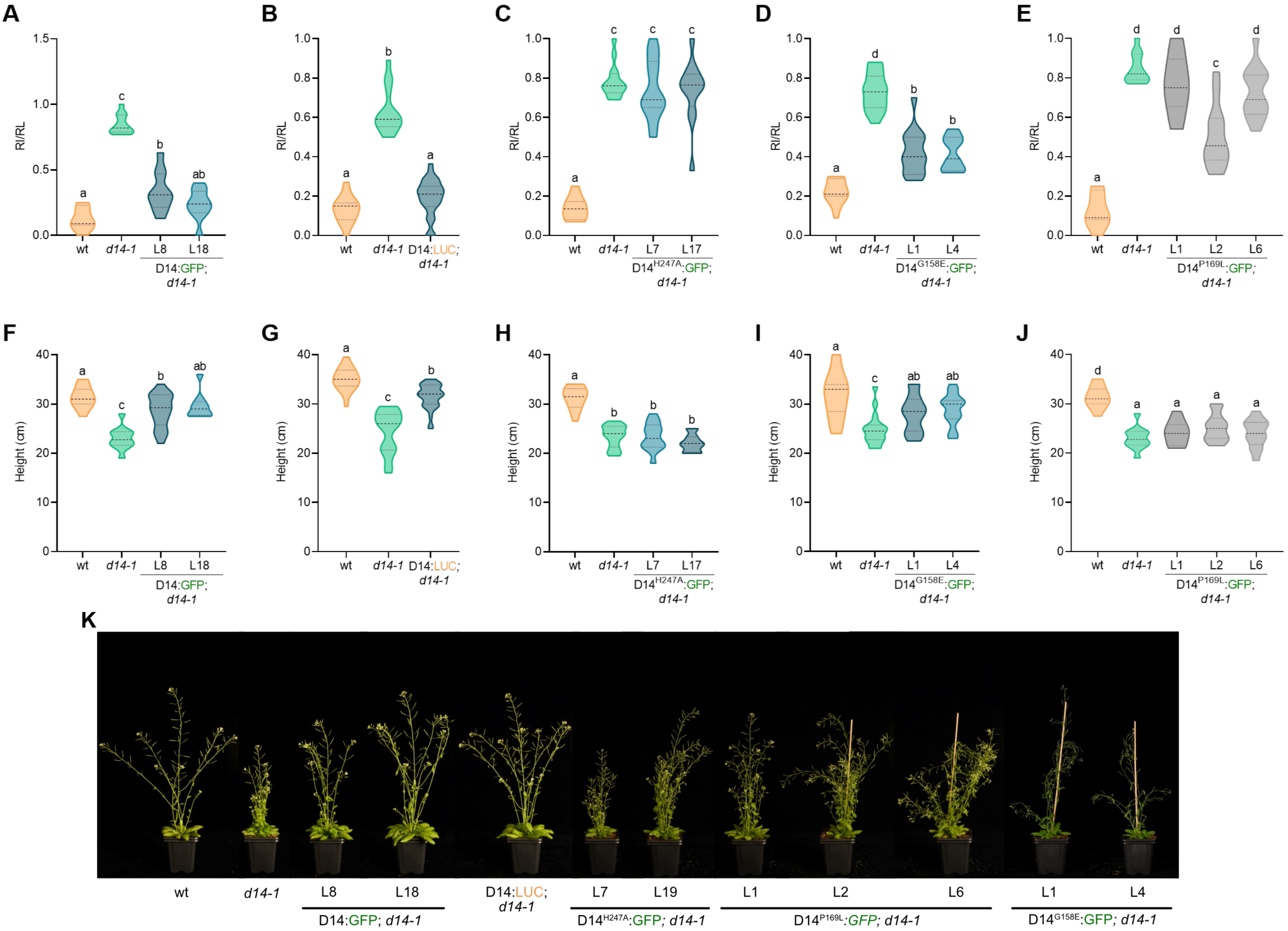


**Supplemental Fig. 1. Functional analysis of D14:GFP, D14:LUC and D14:GFP mutant proteins.** (A-J) Shoot branching (A-E) and height (F-J) phenotypes of wild type (wt), *d14-1* and *35S:D14:GFP;d14-1* lines (A, F), *UB:D14:GFP;d14-1* lines (B, G), *35S:D14*$^{H247A}$*:GFP;d14-1* (C, H), *35S:D14*$^{G158E}$*:GFP;d14-1* (D, I) and *35S:D14*$^{P169L}$*:GFP;d14-1* (E, J) two weeks after bolting. Branching is denoted as number of primary rosette branches/number of rosette leaves (RI/RL). Violin plots represent data distribution. Black dotted lines indicate median values; grey dotted lines delimit the interquartile range (n=11-16). (K) Images of representative individuals from each line. Different letters denote statistical differences in one-way ANOVA with post hoc Tukey test, P < 0.05. For assays with *35S:D14:GFP;d14-1*, line L18 was used.

A

B

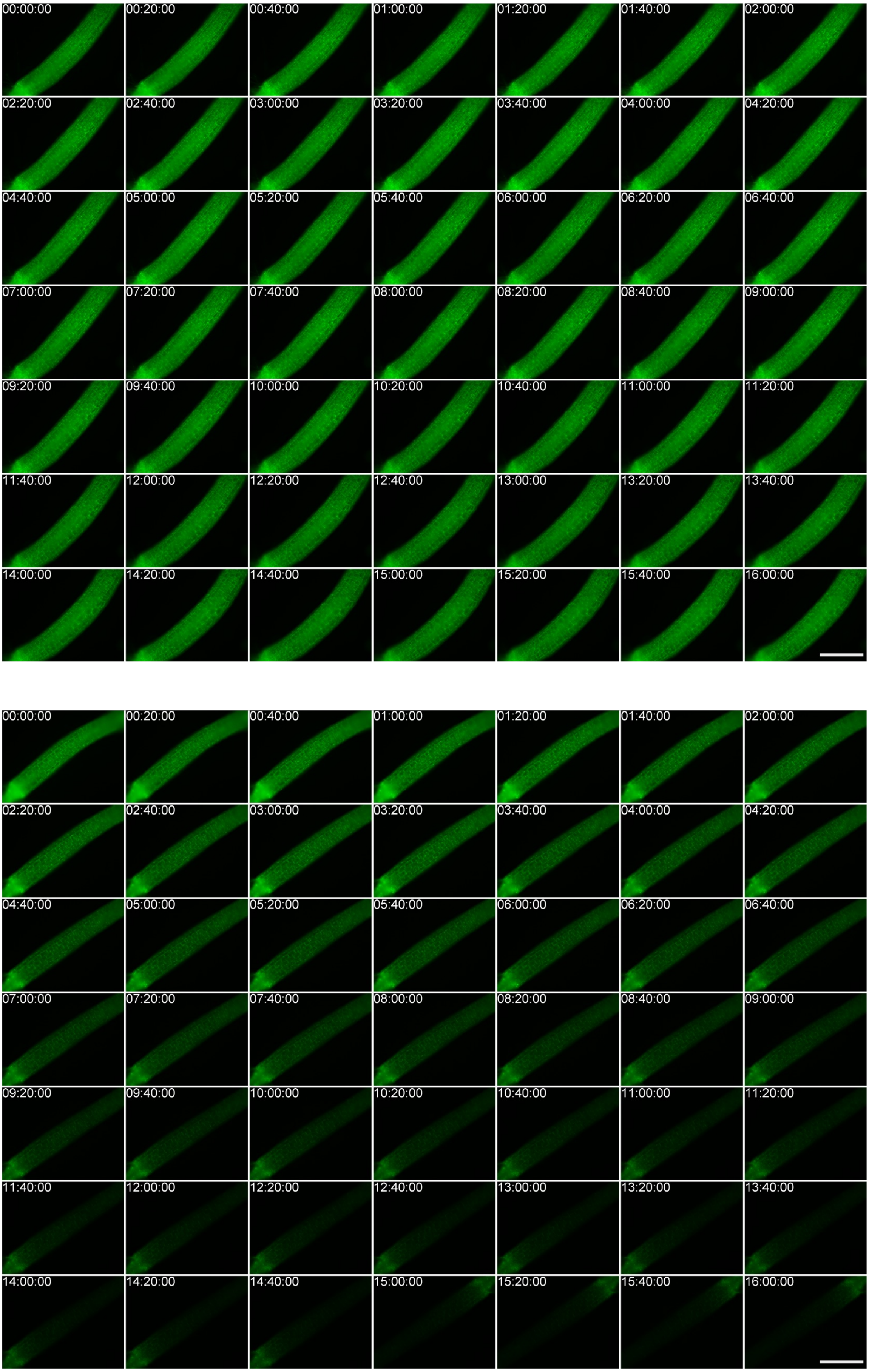


**Supplemental Fig. 2**. **D14:GFP time-lapse assay.** Time series of fluorescence images of the hypocotyl of *35S:D14:GFP;d14-1* seedlings treated with 5 μM $GR24^{5DS}$ (A) or equivalent volumes of acetone as mock treatment (B). Images were taken every 20 minutes up to 16 hours. Scale bar = 500 μm.

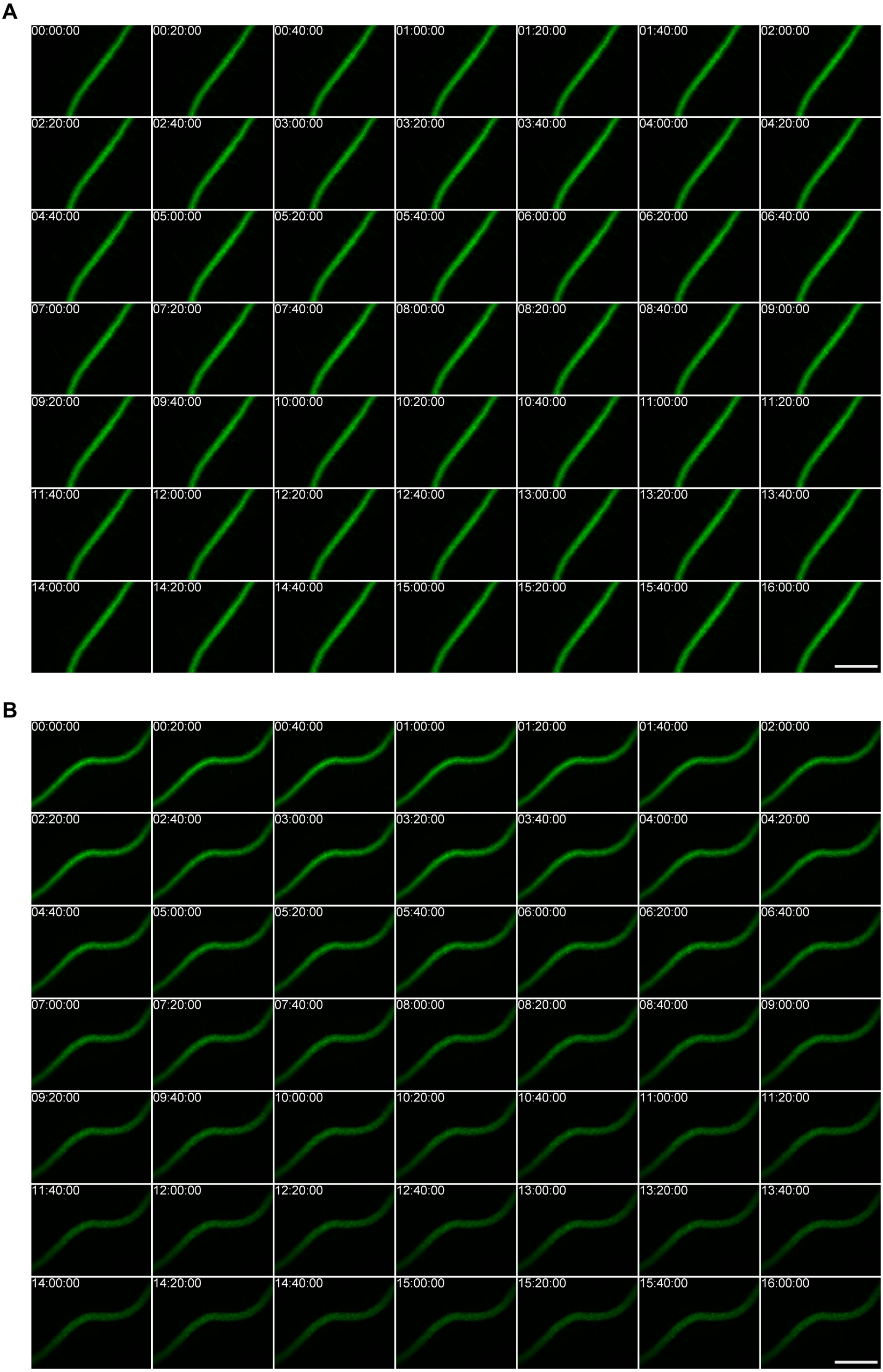


**Supplemental Fig. 3**. **D14:GFP time-lapse assay.** Time series of fluorescence images of the root of the *35S:D14:GFP;d14-1* seedlings treated with 5 μM GR24$^{5DS}$ (A) or equivalent volumes of acetone as mock treatment (B). Images were taken every 20 minutes up to 16 hours. Scale bar = 500 μm.

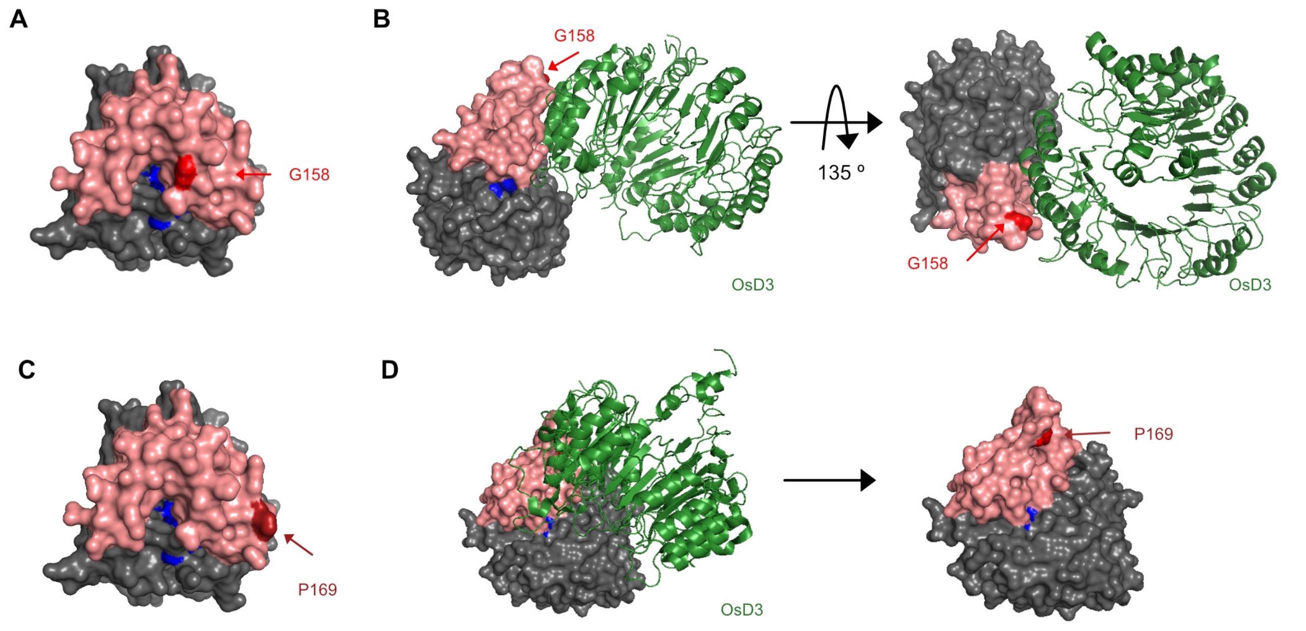


**Supplemental Fig. 4. Localization of the G158 and P169 residues possibly involved in the D14-MAX2 interaction.** 3D molecular visualization as a surface representation of the D14 protein in modified PDB:4IH4 (Zhao et al., 2013) (A,C) and PDB:5HZG (Yao et al., 2016) (B,D). The α/β hydrolase core domain is coloured in grey; the lid domain in salmon; residues of the catalytic triad in blue; residues G158 and P169 are highlighted in red. In B and D, the ribbon representation of the rice MAX2 orthologue OsD3 is coloured in green.

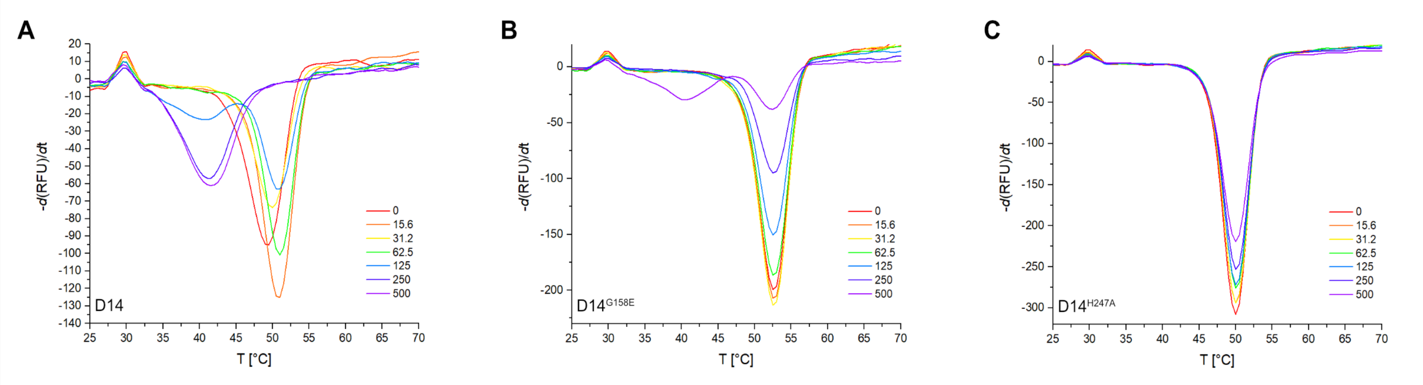


**Supplemental Fig. 5**. **Evaluation of D14-SL interaction and protein stability of D14, D14$^{G158E}$ and D14$^{H247A}$**. In the DSF assays, the melting temperature (Tm) profiles of the D14 (A), D14$^{G158E}$ (B) and D14$^{H247A}$ (C) proteins, each isolated from *E. coli*, were monitored under increasing concentrations of GR24$^{5DS}$, and thermal denaturation shifts observed in these proteins were recorded.

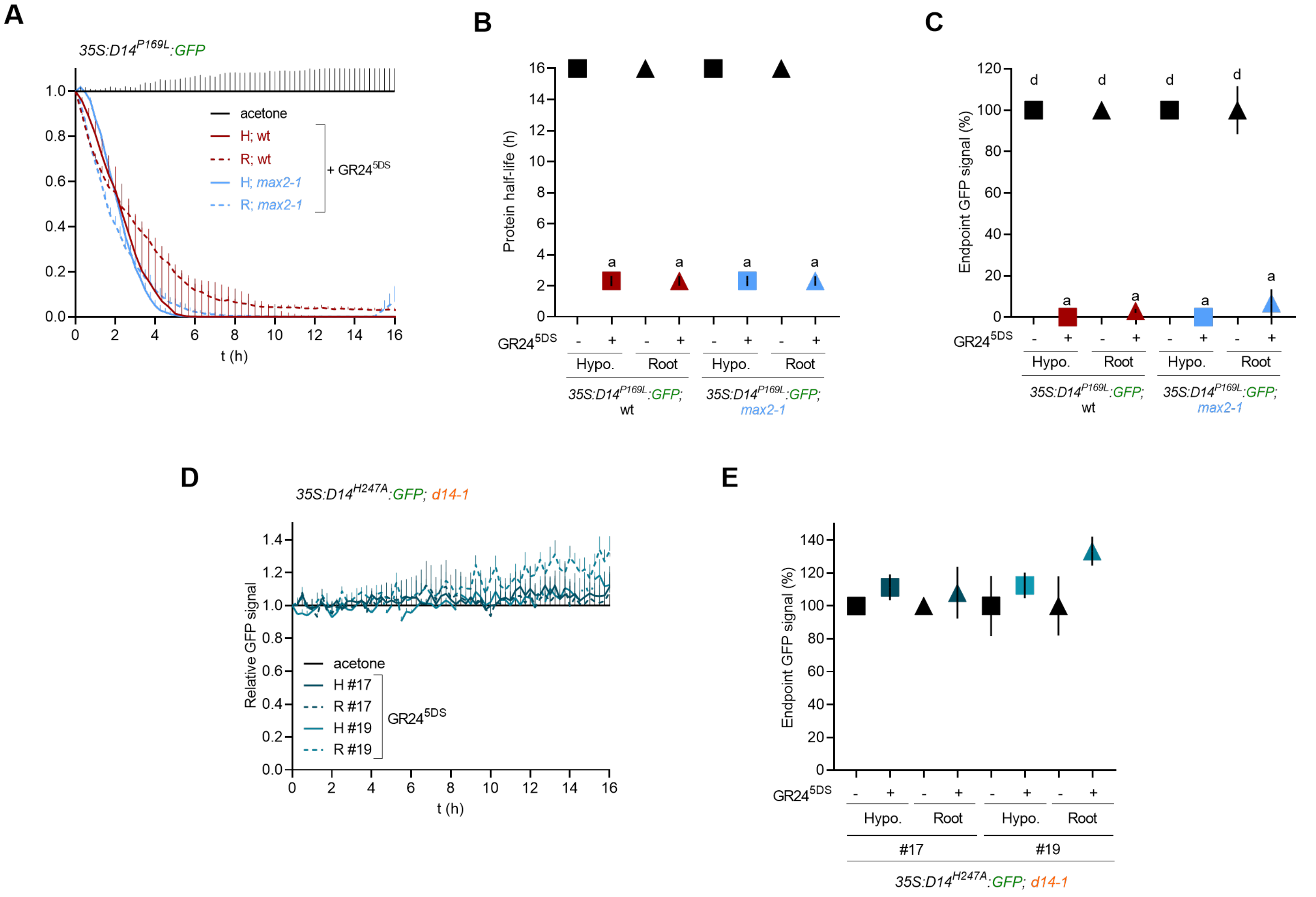


**Supplemental Fig. 6. Analysis of D14$^{H247A}$:GFP and D14$^{G158E}$:GFP stability in response to SL.** (A) Time-lapse fluorescence study of D14$^{P169L}$:GFP signal decay (relative to mock treatment values) of hypocotyls (H) and roots (R) of *35S:D14$^{P169L}$:GFP;d14-1* seedlings treated with 5 µM GR24$^{5DS}$ or acetone (mock). (B-C) D14$^{P169L}$:GFP protein half-life (B) and endpoint signal at 16 h (C) calculated from data in A. Square symbols represent hypocotyls, while triangles stand for roots. (D) Time-lapse fluorescence study of D14$^{H247A}$:GFP signal decay (relative to mock treatment values) of hypocotyls (H) and roots (R) of *35S:D14H247A:GFP;d14-1* seedlings treated with 5 µM GR24$^{5DS}$ or acetone (mock). (E) D14$^{H247A}$:GFP endpoint signal at 16 h calculated from data in D. Square symbols represent hypocotyls; triangles, roots. Data shown as mean ± SE (n=3). In B, symbols clipped at the edge of the Y axis indicate conditions in which protein half-life is longer than 16 h. Different letters denote statistical differences in one-way ANOVA with post hoc Tukey test, $P < 0.05$.
